# Unravelling the effects of disease-associated mutations in TDP-43 protein via molecular dynamics simulation and machine learning

**DOI:** 10.1101/2021.07.28.454112

**Authors:** Abhibhav Sharma, Pinki Dey

## Abstract

Over the last two decades, the pathogenic aggregation of TAR DNA-binding protein 43 (TDP-43) is found to be strongly associated with several fatal neurodegenerative diseases such as amyotrophic lateral sclerosis (ALS) and frontotemporal lobar degeneration (FTD), etc. While the mutations and truncation in TDP-43 protein have been suggested to be responsible for TDP-43 pathogenesis by accelerating the aggregation process, the effects of these mutations on the bio-mechanism of pathologic TDP-43 protein remained poorly understood. Investigating this at the molecular level, we formulized an integrated workflow of molecular dynamic simulation and machine learning models (MD-ML). By performing an extensive structural analysis of three disease-related mutations (i.e. I168A, D169G, and I168A-D169G) in the conserved RNA recognition motifs (RRMs) of TDP-43 and we observed that the I168A-D169G double mutant delineates the highest packing of the protein inner core as compared to the other mutations, which may indicate more stability and higher chances of pathogenesis. Moreover, through our MD-ML workflow, we identified the biological descriptors of TDP-43 which includes the interacting residue pairs and individual protein residues that influence the stability of the protein and could be experimentally evaluated to develop potential therapeutic strategies.

## 1. Introduction

The molecular mechanisms leading to the accumulation of TAR DNA-binding protein 43 (TDP-43) in the central nervous system is a key feature of several common neurological disorders in ageing societies, such as frontotemporal dementia (FTD), Alzheimer’s disease (AD), amyotrophic lateral sclerosis (ALS) and limbic predominant age-related TDP-43 encephalopathy (LATE)^1^. The TDP-43 protein is highly conserved and plays a significant role in RNA regulation such as splicing, transcriptional regulation, mRNA stabilization^2, 3^, etc. Moreover, the TDP-43 is a ubiquitously expressed member of the large heterogeneous nuclear ribonucleoprotein (hnRNP) family that shows specific RNA/DNA binding ability by the highly conserved RNA recognition motifs (RRMs) of the proteins^4^. But during pathological conditions, several post-translational modifications occur in the protein that leads to their cytoplasmic aggregation causing TDP-43 proteinopathies^5–7^. In fact, ~97% of all the cases of ALS, ~75% of patients with severe AD and ~45% of all the cases of FTDL involve the aggregation of TDP-43^8, 9^. And all the four diseases which are together known as TDP-proteinopathies^2, 10^ constitute the major cause of dementia in the world and are expected to rise notoriously in the coming years^11^.

Over the past years, numerous studies were performed to understand the pathological mechanisms underlying TDP-43, and most of the studies focused on the misfolding^8^, mislocalization^12^ and aggregation^13–15^ of the TDP-43 protein. Several studies were able to depict certain mutations in the C-terminal regions of TDP-43 such as R361S, N345K, Q343R that lead to its toxic aggregation in the cytoplasm^1, 8, 16–18^. Certain mutations in TDP-43 such as A382T, A315T, M337V were also reported to escalate its cytoplasmic mislocalization^17, 19^. Most of these mutations are introduced as post-translational modifications, the most common being phosphorylation and ubiquitination of the protein^20^. However, the underlying mechanisms leading to the aggregation of TDP-43 remain elusive. One of the prominent hallmarks of TDP-43 induced proteinopathies is marked by its depletion in the nucleus and increased aggregation into cytoplasmic inclusion bodies. And the rate-limiting step of this process is the cleavage of TDP-43 and generation of C-terminal fragments by the cysteine proteases, caspase and calpain^21, 22^.

The generation of C-terminal fragments, TDP-25 (25kD fragment) and TDP-35 (35kD fragment) by caspase-3 and caspase-7 is the most prominent step for the clearance of TDP-43^23^. Several experimental studies have also reported a significant delay in cell death on blockage of the caspase digestion^23–25^. Moreover, it is shown that out of the four prominent caspase cleavage consensus sites, three sites lie in the RNA recognition motifs (RRM) of the TDP-43 protein^26^. The cleavage at D89-A90 of N-terminal domain (NTD) of TDP-43 generates TDP-35 that is still capable of folding correctly^27^. But, both the cleavage sites at D169-G170 and D174-C175 of the RRM of TDP-43 generate TDP-25 which lacks the NTD, nuclear localization signal (NLS) and most of RRM1, trapping the protein in the cytoplasm^28^ and thus enhancing its cytoplasmic aggregation^29^. Interestingly, certain mutations, particularly D169G in the RRM1 domain is reported to increase the thermal stability of the protein which becomes more accessible to cleavage by caspase-3 resulting in the early onset of diseases such as ALS^26^. They also showed that the neighbouring I168 residue is also very crucial for protein folding^26^. Recently, mutations in the RRMs are also shown to influence the DNA or RNA binding specificities^30^ indicating the role of nucleic acid-binding in TDP-43 aggregation. However, studies on the effect of disease-related mutations on the RRM domains of the TDP-43 is still very limited. Given the crucial role played by the disease-causing mutations in the RRM domain in regulating the protein conformational stability, nucleic acid binding and their role in affecting the cleavage sites remains largely unexplored.

In this paper, we perform extensive structural analysis on the impact of disease-related mutations in the RRM1 domain of the TDP-43 protein. We address this question by formulizing a framework MD-ML, that integrate molecular dynamics and machine learning approaches in mainly two ways (i) structural analysis of the wild type and disease-causing mutant proteins bound to the nucleic acid by molecular dynamics simulations and (ii) identifying the functionally important regions and biological descriptors of TDP-43 in different mutated states that are crucial in explaining the effect of disease-causing mutations in the caspase cleavage sites in RRM motifs of TDP-43 using Machine Learning models. In this direction, the in-silico approach of molecular dynamics simulation holds promising aspects as it provides insights at the molecular level by mimicking the physicochemical changes that occur within the biomolecule when subjected to a specific condition. However, the high dimensional nature of biomolecular simulation data makes it challenging to extract the discriminatory features or set of collective variables (CVs) of the system^31^. In our case, the CVs that we are interested in are those optimum sets of descriptors that discriminate the different biomolecular states (wild type and mutated) over a specific time scale. Recently, machine learning methods are receiving great attention in the biological research domains including sequence structure-function prediction, genomics, and biological imaging^32–34^. Machine learning models have shown esoteric capabilities of learning the ensemble properties through the MD simulation data and subsequently were able to predict the biomolecular functions^35^ as well as the optimum set of CVs or important discriminating molecular descriptors such as long-distance interaction^36^ that govern the biomolecular changes during the trajectory.

## 2. Material and Methods

### 2.1 Molecular Dynamic Simulation

To investigate the structural changes induced by the disease-causing mutants on the nucleic acid bound RRM1 domain of TDP-43 proteins, we performed 100ns long molecular dynamics (MD) simulations of the wild-type and three TDP-43 mutant proteins with D169G, I168A-D169G, I168A mutations in their RRM1 domain. We selected these mutations as they are experimentally found to impact the overall stability of the protein^26^. We adopted the crystal structure of human TDP-43 in complex with a single-stranded DNA (PDB ID: 4Y0F.pdb) for the wild-type protein and crystal structure of human TDP-43 (PDB ID: 4Y00.pdb) for the D169G mutant protein^26^. Due to the unavailability of crystal structures, the I168A-D169G, I168A mutations were introduced into the wild-type protein using the PyMol’s Mutagenesis Wizard. The Isoleucine and Aspartic acid at positions 168 and 169 in the wild type were replaced by Alanine and Glycine respectively in I168A-D169G TDP-43 mutant whereas the I168A TDP-43 mutant is obtained by replacing Isoleucine at position 168 to Alanine. The RRM1 protein used in our study is composed of 78 amino acids and 10 nucleotides. A schematic representation of the protein structures and the position of mutations with respect to the cleavage site is shown in Fig. 1A. All the MD simulations were carried out using the GROMACS software (version 2019.3) and AMBER03 protein, nucleic AMBER94^37^ forcefield. Each wild type and mutant system were solvated using a simple point charge (SPC) water model and were neutralized by adding Na+ and Cl-ions. Each solvated system was subjected to 50,000 steps of energy minimization and 100ps each of NVT and NPT equilibrations. The temperature was kept constant using a Berendsen thermostat and the pressure was fixed at 1 bar using The Parrinello-Rahman algorithm^38^. Further, 100ns long MD simulations were carried out at a mean temperature of 300 K and pH 7 with an integration time step of 2fs. The simulations were carried out under periodic boundary conditions and particle mesh Ewald treated the long-range electrostatic interactions.

**Fig.1:**
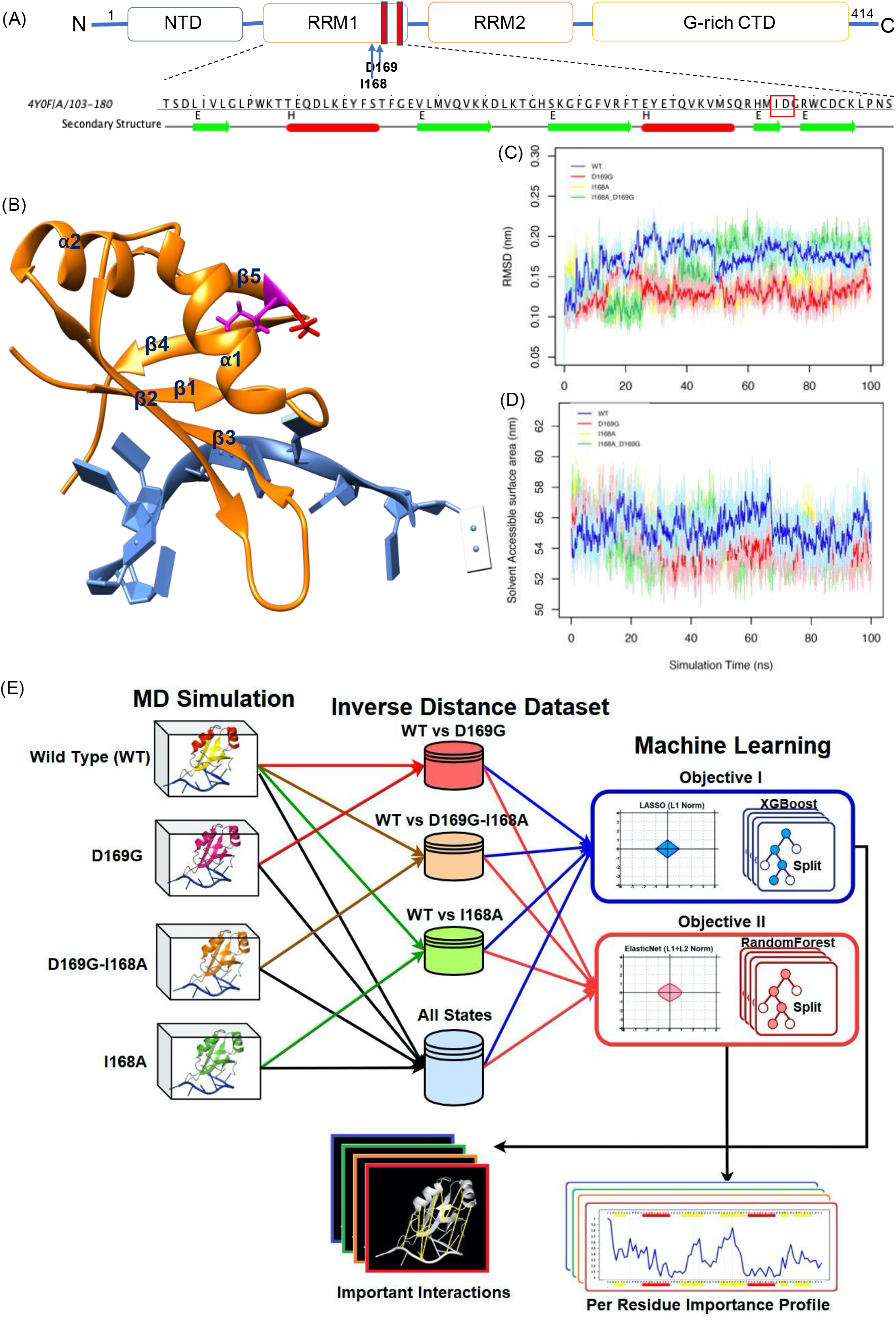
Schematic representation of the DNA bound RRM1 domain of TDP-43 and the MD-ML workflow used for analysing the structural changes in the protein due to mutations. (A) The domain architecture of TDP-43 consists of an N-terminal domain (NTD) followed by two RNA recognition motifs (RRM1 and RRM2) and a Glycine-rich C-terminal domain. The cleavage sites in the RRM1 domain (D169-G170 and D174-C175) are shown in red strips. The sequence of the RRM1 domain is shown with a red box highlighting the positions of the residues to be mutated. (B) The crystal structure of TDP-43 RRM1 domain (PDB ID: 4Y0F.pdb) in complex with a single-stranded DNA. The RRM1 domain consists of two α stands and five β strands. The amino acids that are mutated, i.e., D169G (shown in red), I168A (shown in magenta), I168A-D169G lie in a loop between the β4 and β5 strand and one of the cleavage sites (D174) are in the β5 strand. (C-D) The Root Mean Square Deviation (RMSD) and Solvent Accessible Surface Area (SASA) of the WT and mutant proteins (D169G, I168A, I168A-D169G) are shown as a function of the simulation time(ns). (E) Schematic representation of the MD-ML workflow. The implementation of Machine Learning algorithms over the Molecular Dynamics simulation data to interpret significant biological characteristics.

The analyses were performed using the GROMACS utilities and the coordinate dataset generated through the MD simulation was used to generate (i) the average contact-map for the entire trajectory of each state (wild type and mutants) where the contact between every i^th^ and i+4^th^ residues were calculated with a cutoff distance of 10 Å. (ii) difference of the average pairwise distances of the residues in the mutant proteins with respect to the WT protein and (iii) Dynamic cross-correlation (DCC) analysis was carried out to intuitively unravel the functionally relevant regions of protein states throughout the trajectory. Within the given molecule the DCC metric for a pair of residues represents the degree to which these residues move together and is calculated as^39^:

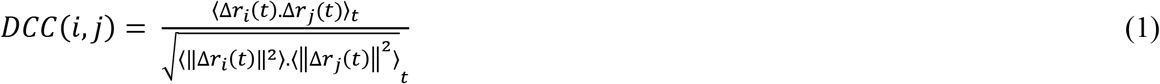

where r_i_(t) is the vector of the i^th^ atom’s coordinates as a function of time t, 〈*〉 denotes the time ensemble average and Δ*r_i_*(*t*) = *r_i_*(*t*) — 〈*r_i_*(*t*)〉_*t*_. Although the contact-map and DCC approach highlighted some ambiguous yet important fingerprints of each state, these approaches however provided little information to unmask the intricate intra- and inter-molecular relations and features that are critical for understanding the effect of different types of mutations in the protein.

### 2.2 The Machine Learning Utilization

The dataset generated by the MD simulation is high dimensional, extraneously noisy and encompasses a set of important features and patterns encrypted within that could explain the changes that occur within the biomolecular ensemble. Machine learning models carry promising potentials to identify these discriminatory features and fingerprints that could elevate our understanding of the key mechanisms underlying the behaviour of biomolecules in the specific state.

#### 2.2.1 Data Pre-processing

There are several ways in which the MD simulation dataset can be utilized to extract features. For a large system, the internal coordinates of residues could be used as feature variables but often tend to produce fallible conclusions due to overfitting^36^. For a small or moderate size ensemble, such as the TDP-43 DNA complex in our case, the distance-based features would provide much deeper insights as it contains a profuse amount of conformational and physicochemical information of the system^36^. Here we incorporated the inverse inter-atomic distance as features because it also highlights the significant local changes within the ensemble. For amino acid, the C_α_ and for nucleotides, the centre of mass was used to measure the inverse inter-residue distance. The total of 88 residues including that of the RRM1 domain and the nucleotides resulted in 88(88-1)/2 variables. The 2000 frames generated for each type of state thus yield an inverse-distance dataset of size 2000 x 3828. To meticulously probe the effects of the mutations, we adopted two approaches (Fig. 1E) i.e. (i) To study the individual effects of the point mutations on TDP43, we performed the feature extraction by pairwise concatenation of the datasets of each type of mutated state (i.e. D168G, D168G-I169A and I169A) with the wildtype (WT) dataset; (ii) To unmask the globally important interactions and discriminatory CVs, we concatenated the labelled inverse-distance dataset of all the mutated and wild type states. To subdue the curse of dimensionality, we included only those interaction pairs as variables whose interatomic distance was greater than 10Å but also have a distance less than 10 Å in at least one frame.

#### 2.2.2 Machine Learning models

Here, we employed supervised machine learning models over the simulation datasets - (Objective I) to extract the highly discriminatory features (i.e., interacting pairs) that distinguish the mutated states from the wildtype state of TDP43 and its binding; (Objective II) to depict the importance per residue profile of TDP-43 that unmasked the biological descriptors at the residue level. For the former, we employ two feature extraction models L1 regularization regression model (LASSO) and Extreme Gradient Boosting Machine (XGBoost). For the latter, we incorporate L1-L2 regularization regression model (Elastic Net) and Random Forest (RF) models (Fig. 1E). Based on their scaled scoring scheme, these models would extract important pairs of interactions but the relevance per residue was computed by averaging the importance score of all the pairs involving that residue followed by normalization between 0 and 1. Models were trained and tested over the concatenated dataset where each frame was labelled to its corresponding state. For feature selection, we only corroborate the supervised learning model because, unlike unsupervised models, supervised learning models are known for their esoteric capabilities of exploiting the class information to learn the encrypted discriminatory features and subtle patterns within the dataset that highlights the key differences between given classes.

##### 2.2.2.1 Regularized Regression Models

###### Least Absolute Shrinkage and Selection Operator

(LASSO) is a regularized regression model from the family of generalized linear models. By penalizing the (L1-norm) regression model, LASSO reduces the regression coefficient to zero for those features that have low contribution in the learning of the model. This makes LASSO a pervasive feature selection model. Alternative to LASSO regression, Ridge regression is also based on the model penalization, but it involves an L2-norm^40^.

For a given feature dataset 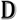, let *x_ij_* be the observation of the *j^th^* variable (1 < *j* ≤ *p*) in the *i^th^* (1 < *i* ≤ *N*) frame and consider *y_i_* be the corresponding state label of the *i^th^* frame. In addition to minimizing the sum of squared error, the regularized model learns the regression coefficient *β_j_* for each *j^th^* variable including the intercept *β*_0_ (eq.2) by imposing a constraint on the coefficient *∑J*(*β_k_*) ≤ *t*^41^.

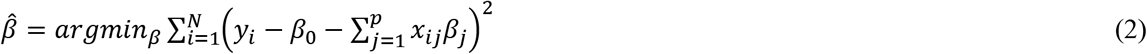

LASSO truncates the coefficient of the low-importance to zero features by imposing L1 constraint (*J*(*β_k_*) = |*β_k_*|) (eq. 3) while the Ridge shrinks the coefficient close to zero for the low contributing variable by imposing L2 constrain 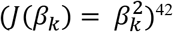^42^.

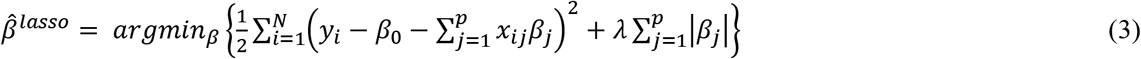

where *λ* is the penalty parameter and we find the best fitting *λ* using cross-validation^40^. While LASSO identifies a set of most discriminating independent inter-residual interaction pairs, it drops all the other non-contributing as well as low-contributing variables and therefore is not suitable to account for per-residue importance for the system. We, therefore, employ an Elastic Net classifier to yield a per-residue importance profile for the mutated systems. Based on penalization, Elastic Net is a smart combination of L1 and L2 constrain with *J*(*β_k_*) (coefficient constrain) given as:

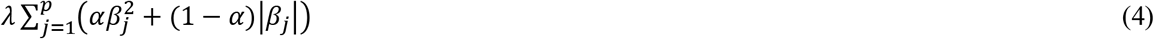

where the *α* constant governs the intensity of Ridge and LASSO penalties^41^. We used the R package “caret” to train LASSO and Elastic Net models^43^.

##### 2.2.2.2 RandomForest

Breiman et. al^44^ developed the random forest (RF) algorithm that utilizes the ensemble of decision trees to perform prediction and classification. After bootstrapping the feature dataset with replacement, allowing duplicate entry of the instances, the fully grown classification trees are produced by randomly sampling a set of variables at each split^45^. This way, RF performs feature selection by carrying out a bootstrapping and aggregation (bagging) for tree building. The decisions are then made based on the average of collective decisions by all the fully grown decision trees. For each generated tree, the RF quantifies the performance of that tree by measuring Out-Of-Bag (OOB) error based on the samples that were not included during the bootstrapping. The OOB has a crucial role in estimating the goodness of fit for RF while also exempting us from performing the cross validation. For a decision tree, the Gini impurity metric reflects how good a node is in splitting. In RF, the mean decrease in impurity for an individual variable over all the trees indicates the importance of that feature variable^44^. We used the R package “RandomForest” to train RF model^46^. A detailed description of the RF algorithm is included in the supplementary text.

##### 2.2.2.3 XGBoost

J.H Freidman^47^ introduced gradient boosting algorithms that have shown competitive classification and prediction potentials on many occasions. Based on the gradient boosting algorithm, Chen et al.^48^ proposed a scalable tree boosting machine called extreme gradient boost (XGBoost) that performed remarkably in several biological research^49, 50^.

The underlying principle of XGBoost is that for the given data 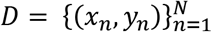, the *k* classification trees or stumps given as *F* = {*f*_1_(*x*), *f*_2_(*x*)…, *f_k_*(*x*) } assigns a class decision score *f_k_* to the data instance by passing it through the leaf node as per the division points of the variable. When the data instance *x_i_* passes through a leaf node, the *f_k_* is assigned and the prediction result is registered. The collective sum of prediction results by each tree accounts for the final prediction result for each instance. The model follows:

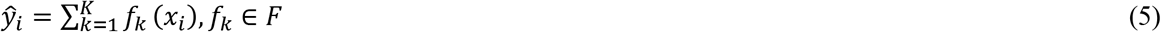

Here, *f_k_*(*x_i_*) denotes the prediction score of *k^th^* leaf node. The sum of prediction results overall *κ* stumps yields the prediction result (*y_i_*) for the *i^th^* instance *x_i_*. The Objective function *Obj*(*θ*) for XGBoost is given as:

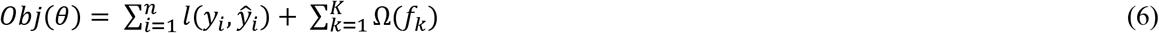

Where θ is the model parameter. The sum 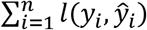 accounts for the error that the model accumulates with *l*(*x*,*y*) is the error function. The second summation 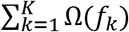 represents the model’s regularization term, accounting for the complexity of the trees. A thorough overview of the XGboost algorithm is described in Wei li et al^51^. An inclusive pseudocode^47^ for the gradient boosting algorithm is added in the supplementary text. Here we employed XGBoost to identify the most influencing residue interaction pairs for the mutated states. We used an R package “XGBoost” to implement the algorithm^52^.

##### 2.2.2.4 Principal Component Analysis (PCA)

In the paradigm of finding biological descriptors using a simulation dataset, the unsupervised learning models perform worse than the supervised learning model because of their incapability to exploit the state label^36^. While the unsupervised learning models are not suitable for the unmasking of descriptors, they are highly robust in dimension reduction where the feature variables are mapped to a lower-dimensional configurational space and thus highlighting the main characteristics of the trajectory. In this direction, the PCA method is the most celebrated machine learning model that reveals the dominant modes in the motion of biomolecules by incorporating the molecular dynamic simulation trajectory^53, 54^. These motions reflect the correlated vibrations or collective motion of a set of residues in the trajectory^53^. In principle, For the N residue system, the 3-dimensional cartesian coordinate configuration is transformed into a 3N × 3N covariance matrix whose eigenvectors provide vectorial information indicating the direction of each component of the trajectory while the corresponding eigenvalue represents the intensity contribution of that particular component. We elucidated this motion of eigenvectors through “porcupine” plots^55^ for each type of state of TDP43 which depicts the direction and magnitude for each residue in that state. We used PyMol software to generate the porcupine plots^56^.

The computation was performed on an Intel (R) Core (TM) i5-4310 U, 16 GB RAM, and 64-bit OS Win 10 configuration. The R version 4.0.3. was used to prepare codes for conducting the experiments and is shared in the Github link.

## 3. Results and Discussions

### 3.1. Molecular insights into structural changes in protein due to the disease-causing mutations

To investigate the structural changes that undergo in the protein due to mutations, we started by analyzing the stability of the single-stranded DNA bound wild-type (WT) and the mutant proteins from the 100ns simulation trajectories. From the RMSD analysis of the proteins (Fig.1C), we see that the overall protein structure is maintained for the wild type and mutant proteins and the mutations in the protein don’t impact the overall conformational stability of the protein molecule. However, we see that the D169G mutant protein shows the lowest RMSD as compared to the wild-type protein (Fig.1C). Several experimental studies have indicated that the D169G mutation increases the overall stability of the protein due to enhanced packing or hydrophobic folding of the RRM1 core domain^26^. We also calculated the evolution of solvent accessible surface area (SASA) of the WT and mutant proteins throughout the simulation trajectory and found that mutant proteins, especially the I168A-D169G and D169G mutation has a lower SASA indicating stronger hydrophobic interactions of the core residues of RRM1 domain (Fig.1D).

To further investigate the underlying mutation-induced molecular changes in the protein, we estimated the structural changes occurring in the WT and mutant proteins throughout the simulation time. Firstly, we calculated the difference of the average pairwise distances of the residues in the mutant proteins with respect to the WT protein to view the relative structural changes occurring in the protein due to mutations. The averaging is done over an ensemble of configurations of WT and mutant proteins generated throughout our simulations. From Fig. 2(A, C, E), we see that some of the inter-residue distances increase in the mutant proteins with a simultaneous decrease in other inter-residue distances. Although the global dissemination is observed for all the mutant protein structures, the major impact is seen in the double mutant protein (Fig. 2C) where we see that the relative distances of the start (α1 and β1 strand) and end regions (β4 and β5 strands) of the protein decreases with respect to the mutant protein core (β2, β3 strands). A similar but less profound observation is seen for the D169G mutant protein (Fig. 2A). The relative motion of the mutant proteins with respect to the protein core is shown in the porcupine plots in Fig. 2 (B, D, F). The porcupine plots show the coordinated motion of the C_α_ atoms in the mutant proteins along their first eigenvectors. The arrowhead shows the direction of motion, and the size of the arrows gives the magnitude of the motion. The porcupine plots further strengthen the observation of the induced movement of the protein terminal regions towards the protein core due to the mutations. After having seen the relative changes in the protein structure due to the mutations, we also analysed if these structural changes in the mutant proteins result in any changes in the contact form within the mutant protein structures throughout the simulation time (Fig. S1A-D). To get a more comprehensive outlook from our non-bonded contact formation map, we also included how long a contact is formed between two residues in our analysis. The darker the spot in the contact map, the longer the contact was formed between those pairs of residues. We see that the overall contact formation in the WT and mutant protein structures remains intact but some new contact formations are observed between the α1 and β5 strands in the I168A-D169G and I168A mutant proteins. This may be due to the removal of the bulkier isoleucine by alanine that facilitates the protein to attain a more compact structure by promoting contact formation between the α1 and β5 strand, mainly between the residues Lys120 – Cys173; Tyr123 – Cys173 and Phe124 – Cys173. This also highlights the interactions of Cys173 as crucial to maintaining the increased hydrophobic interactions of the protein inner core. Several biochemical studies have identified the Cys173 to promote TDP-43 cross-linking via disulphide bond formation leading to decreased solubility of the protein^23^. We further investigated whether the structural changes in the protein-induced due to the mutations affect the relative motions of the protein strands with respect to the protein core. For this, we performed the dynamic cross-correlation analysis of the C_α_ pairs of WT and mutant proteins throughout the simulation trajectory. From Fig. S2(A-D), we see that the protein core is involved in anticorrelated motion with the terminal regions of both the WT and the mutant proteins, whereas significant changes in the overall correlated motion of the protein molecule due to the introduction of mutations were not found. Recent experimental evidence is available substantiating that the D169G mutation in the protein facilitates the hydrophobic inner core of the protein and thus makes it more stable than the WT protein^26, 57^. Our study reveals that along with D169, I168 is also very crucial in maintaining the protein conformation and the I168A-D169G double mutant facilitates a tighter protein core, orchestrating increased stability and thus, inducing pathogenesis.

**Fig. 2.**
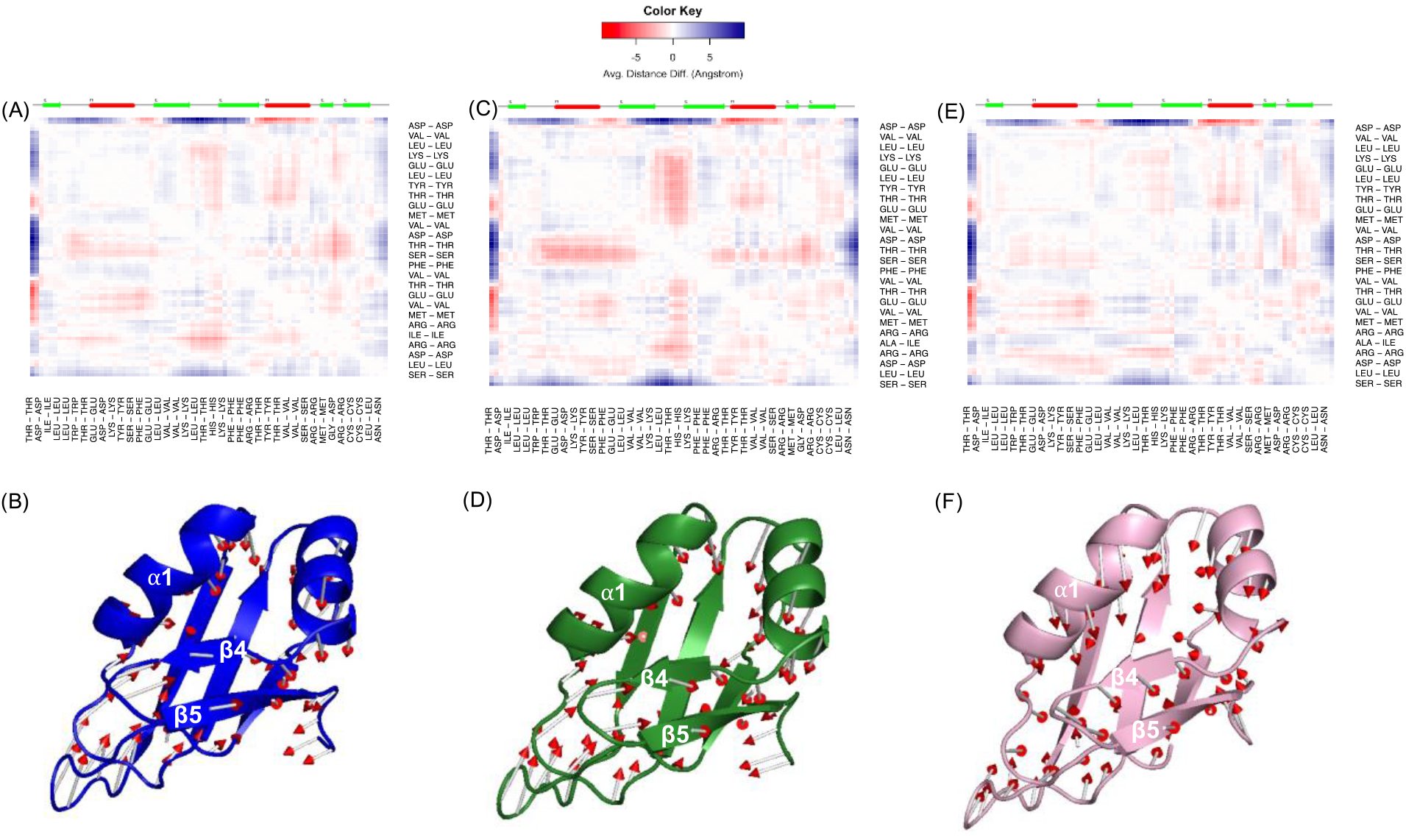
The difference distance map of the (A) D169G (C) I168A-D169G and (E) I168A mutant proteins with respect to the wild-type protein structure throughout the simulation time. The above bar represents the secondary structure of the protein in 2D format; green regions represent beta sheets and red regions represent alpha helix. For clear representation, every third residue in the protein sequence is labelled in the plots. Porcupine plot of (B) D169G (D) I168A-D169G and (F) I168A mutant proteins generated from the first eigenvectors showing the direction of motion of the protein represented by the red arrowheads.

### 3.2 Effect of mutations on interactions of the protein with DNA

Studies have suggested that the disruption of RNA or DNA binding to the RRM1 domain of TDP-43 through mutation or truncation can alter the protein solubility leading to the appearance of aggregates in the nucleus^30, 58^. To investigate the effects of mutation on the interaction of the RRM1 domain with the DNA molecule, we calculated the evolution of distance between the WT and mutant proteins with respect to the DNA molecule. Fig. 3 presents the evolution of the average pair-wise distance between the WT and mutant proteins with the DNA throughout the simulation time. We see that the average residual distance between the core (mainly β4 and β5) and the terminal region (α1 and β1) of I168A-D169G double mutant protein increases the most as evident from the colour-code representing the higher average residual distance between them (Fig. 3C). We also calculated the protein-DNA contact formation in the WT and mutant protein throughout the simulation time and find that most of the contact form with the DNA takes place between the protein core and initial alpha-helix and beta-strand regions of the protein (Fig. S3A-D). From the propensity of contact formation, we see that the introduction of mutations, especially the I168A leads to the disruption of contacts between the protein core, mainly loop4 (between β2 and β3) and the DNA.

**Fig. 3.**
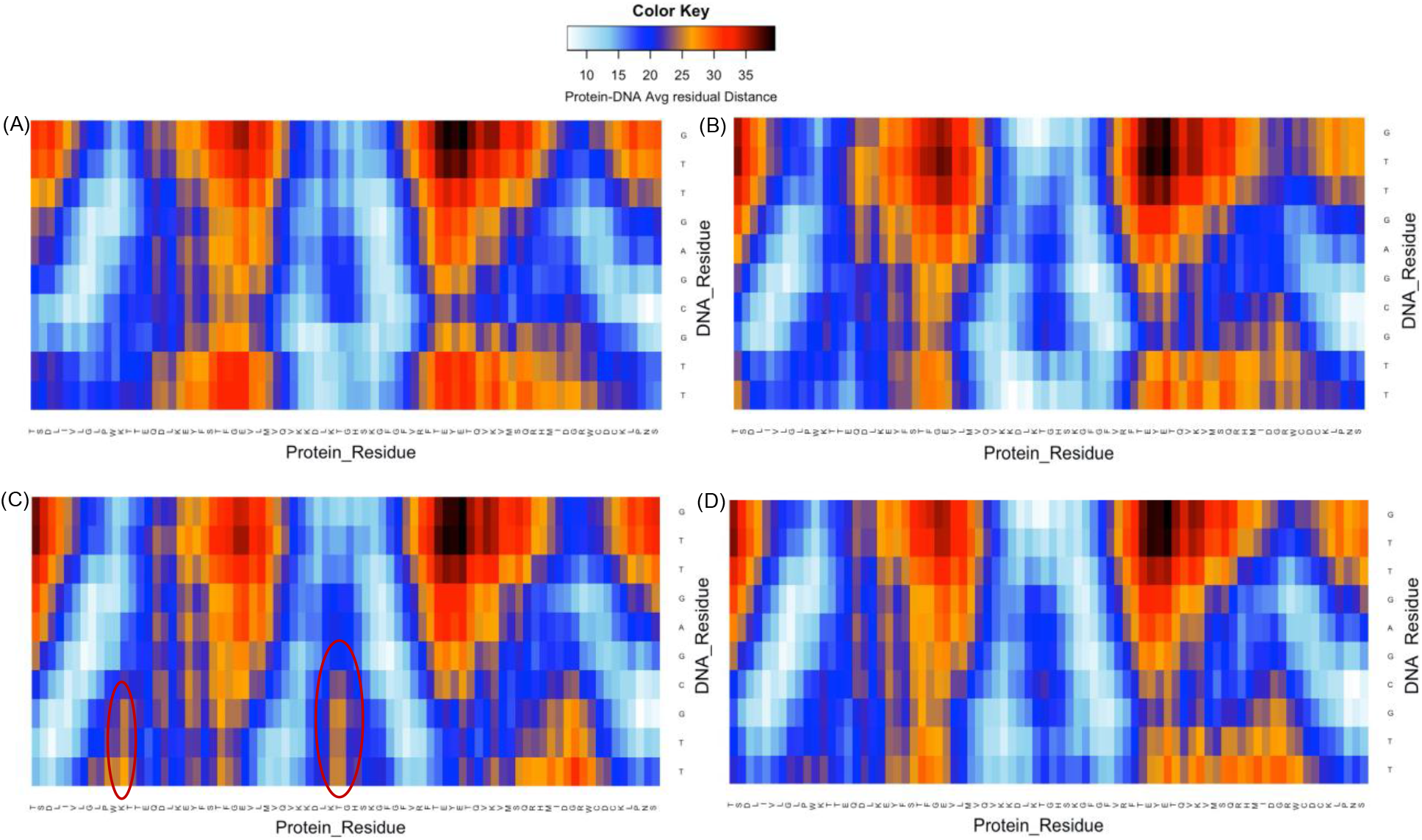
Average residual distance between the (A) Wild-type (B) D169G mutant (C) I168A-D169G and (D) I168A mutant proteins with the DNA throughout the simulation time. The colour-code represents the evolution of the average residual protein-DNA distance from 0 to 50Å. The protein and DNA residues are presented by their single-letter codes.

### 3.3 Identifying the biological important descriptors of TDP-43 in different mutated states

After having seen the structural changes occurring in the protein due to the disease-associated mutations, we now investigate the biological descriptors and the functionally important regions of the protein using Machine Learning models. In this direction, we trained various state-of-the-art supervised learning models to classify the states by learning the ensemble’s features. Interestingly, all the supervised learning models have performed excellently in learning the differences among different states of mutation as they attain a high classification accuracy of 99.9% on a 5-fold cross-validation test (Table S1). The receiver operating curve (ROC) and area under it (AUC) remained very high as well (Fig. S4), thus empowering the results obtained. The list of the most optimum hyperparameters for each type of learning model that were used is provided in the supplementary (Table S2). While probing all the four types of states together, the regularized regression models (LASSO and ElasticNet) remained time expensive for both objectives. This was also observed when conducting a pair-wise WT vs Mutant state analysis.

#### 3.3.1 Objective I – Identify the discriminatory features of the RRM1 domain of TDP-43

To identify the discriminatory features, we employ LASSO and XGBoost on the labelled frame of feature datasets. The importance of each feature is yielded by averaging the importance for the interacting pairs obtained by the models. Figure 4A & 4C presents the average globally important protein inter-residue and protein-DNA residue interaction map respectively where the importance of interaction is shown by the colour key, where the darker shade represents the higher importance of the interaction. Here, with “globally” we tend to mean that the analysis was carried out by training the features of all the mutant states together. The importance interaction map for the pair-wise analysis of each mutant state is provided in the supplementary (Fig. S5). Some of the important protein-protein and protein-DNA interactions are shown in the cartoon representations (Fig. 4B & 4D) of the protein complex. For the protein, we find that most of the important discriminating interactions involve the residues in the protein core (Fig. 4A). The residue pairs having the higher important discriminating scores were found to be M132 - V135, L120 - M132, L111-K136, D119-V133. These interactions can be experimentally assessed by biomedical researchers to develop potential therapeutics for the TDP-43 associated diseases. This also indicates the crucial role played by the protein’s inner core in maintaining the stability of the protein and its interaction with the DNA. These results obtained through the machine learning models are in line with the studies that have suggested that the D169G mutant enhances the stability of TDP-43 by increasing the hydrophobicity or compactness of the core of the RRM1 domain.

**Fig.4.**
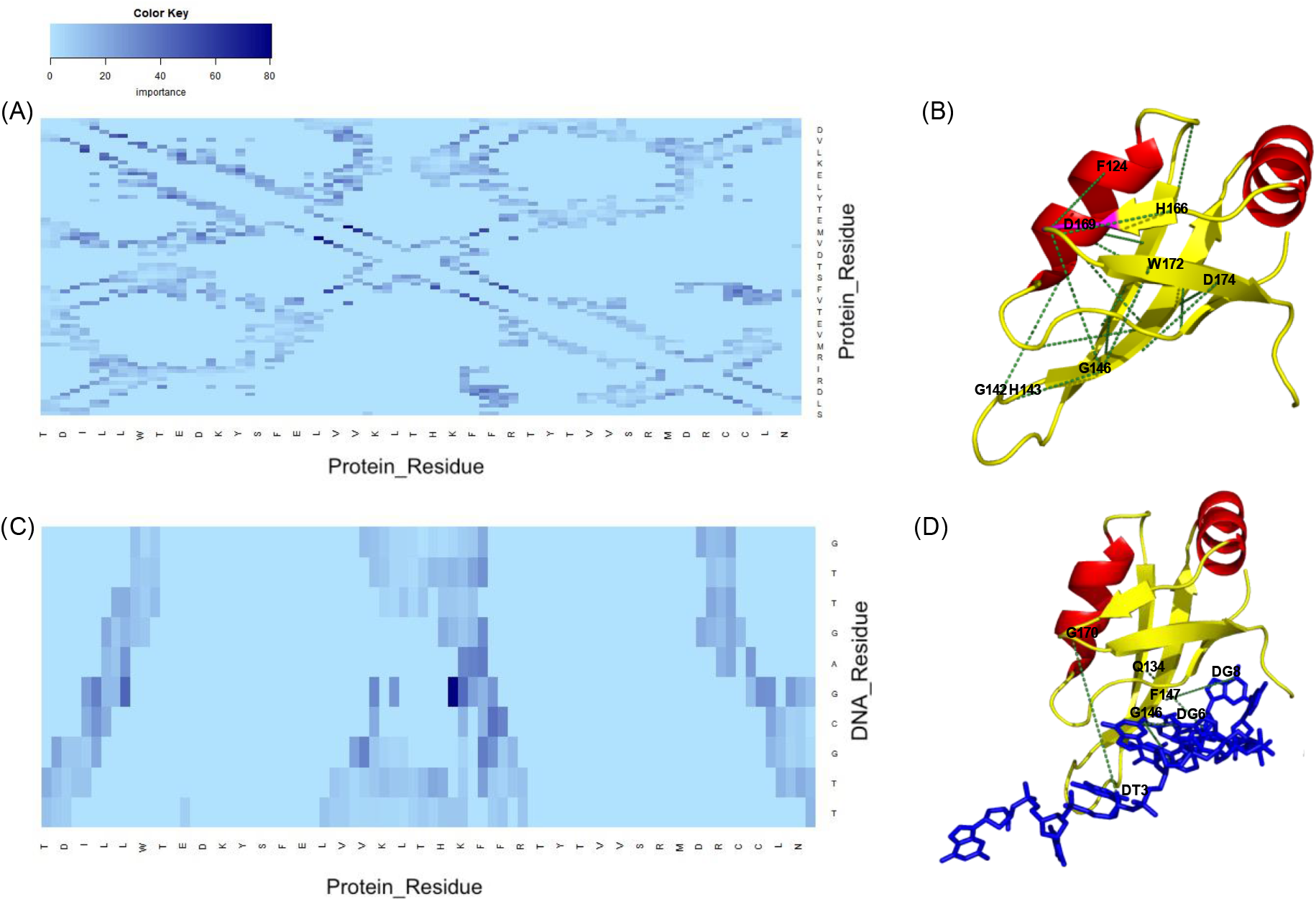
Global average Importance Interaction map of (A) Protein-protein interactions and (C) protein-DNA interactions. The colour intensity shows the importance of the interaction where the darker shade represents higher importance. For clear representation, every third residue in the protein sequence is labelled in the plots. Schematic representation of some of the important (B) protein-protein interactions and (D) protein-DNA interactions. The residues are represented in their single-letter amino acid code. The dashed lines indicate the interacting residue pairs.

In Fig. 4B, we have schematically shown some of the important interactions which involve the mutation site, D169 and the W172 and D174 residues in the β5 strand of the protein where the protein is generally cleaved. We see that D169 residue has important interactions with the residues of the protein core such as G146 (shown in the diagram), W123, and also has important discriminatory interactions with the residues of loop 1 (between α1 helix and β1 strand) such as P112. Moreover, both the mutation site residue, D169 and I168 are found to have important discriminatory interactions with T115 which is also experimentally validated to be crucial in maintaining the protein stability^26^. The cleavage-prone residues in the β5 strand are found to have important interactions with the protein core residues, such as W172 - G146, F147 - D174, F149 - D174, V161 - D174. The importance of the core residues can also be seen in the interaction of the protein with the DNA where the core residues such as Q134, F147, G146 is found to have an important interaction with the DNA molecule.

#### 3.3.2 Objective II-Identifying individual important residues in the RRM1 domain of TDP-43

While unravelling the important interacting pairs, it is also necessary to identify the contribution of the individual residues of the RRM1 domain that plays a crucial role in maintaining the stability of the protein. For a given residue, its importance is calculated by averaging the importance of all the identified interaction pairs that include that particular residue. Since LASSO and XGBoost drop all the moderate to low contributing interactions, they are abysmal to depict the true per residue importance profile. Consequently, the ElasticNet and RandomForest models were employed. Interestingly, the per residue importance profile indicated that the protein core comprised of the β3 and β4 strand of the RRM1 motif shows the highest importance per residue (Fig. 5A). The highest importance per residue in the β3 and β4 strands are found for the residues Q134 and G148. Among the top 15 highest important residues (marked by dashed lines in Fig. 5A) include the I166 and I168 residues of β4 and residues spanning the β1 and α1 strand.

**Fig. 5.**
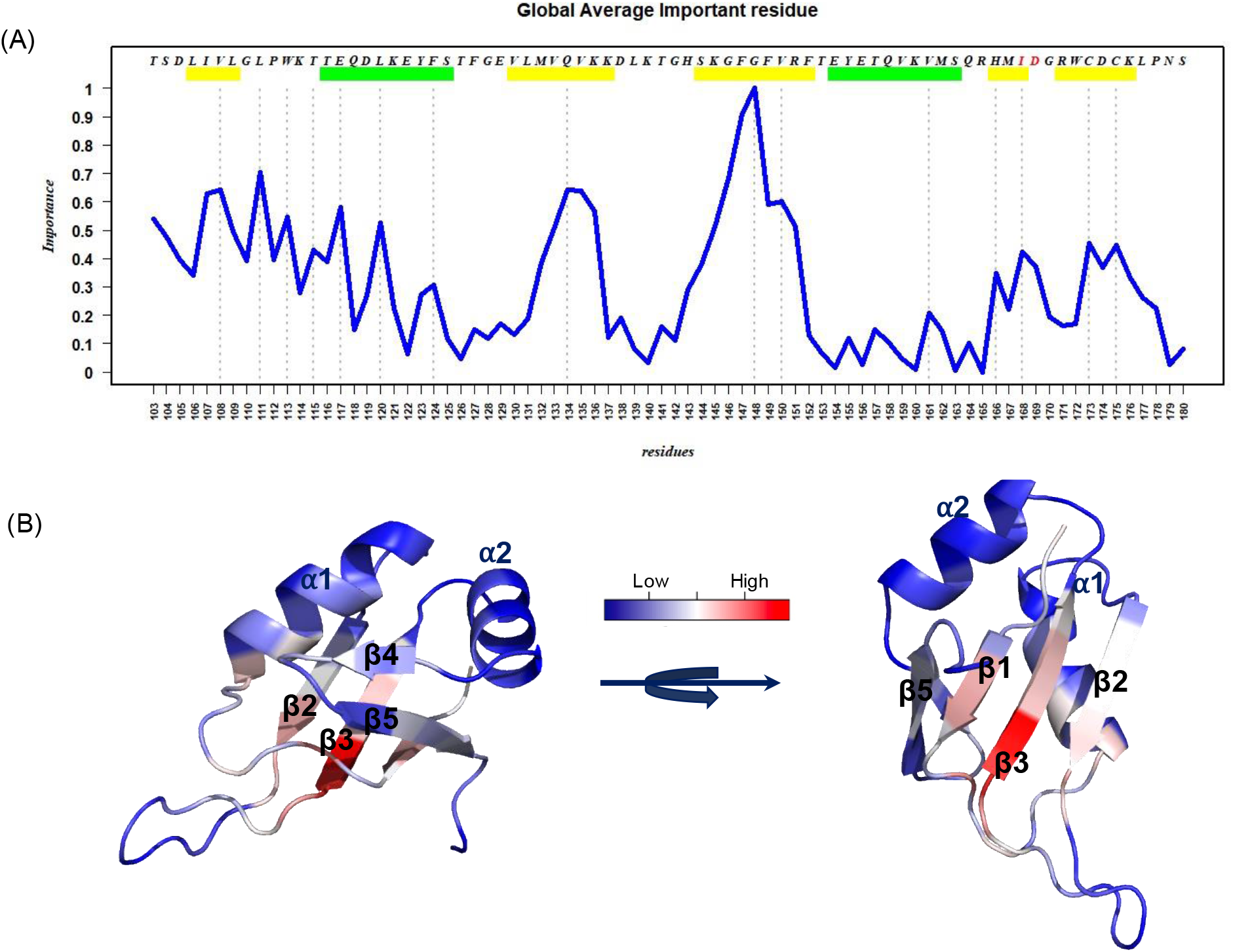
(A) Global average Important residue map of all the residues in the RRM1 motif of TDP-43 protein. The green and yellow bars represent the alpha-helix and beta-strand respectively. (B) Gradient colour coded important residue cartoon representation of the RRM1 protein. The redder shade represents higher importance and the transition from red to blue indicates a decrease in the importance scores.

Fig. 5B shows a schematic representation of the residue importance where the redder shade represents higher importance, and the blue shade represents less importance. This also suggests that the residues in the α1 strand, especially E117 and L120 shown by lighter blue shade are as important as the mutant residues and can also influence the protein stability, which can be experimentally tested by mutagenesis studies. Although similar observation is seen for the interaction map of the individual mutant proteins [Fig. S6], the per residue importance of the protein core region is highest for the I168A-D169G double mutant as compared to the other two mutations. Hence, our analysis not only elucidates the underlying molecular basis of disease-associated mutations in the TDP-43 protein but also helps in identifying the crucial residues involved in the protein stability which can be genetically engineered to address its pathogenesis.

## 4. Conclusion

Recently, neurodegenerative disorders linking to TDP-43 malfunction have increased considerably and mutational modifications in the protein are found to expedite its toxic cytoplasmic aggregation^8^. In this work, we performed an extensive structural analysis on the effect of common disease-causing mutations on the RRM1 domain of TDP-43 protein combining molecular dynamics simulations with machine learning models through the MD-ML workflow. Out of the three mutations studied, D169G, I168A-D169G and 1168A, we found that the I168A-D169G double mutant shows the highest packing of the protein inner core, indicating more stability and hence can lead to the enhanced level of pathogenesis which needs to be experimentally validated in the future. Moreover, using machine learning approaches, we identified the important (i) protein-protein and protein-DNA interacting pairs and (ii) individual protein residues that are crucial in maintaining the stability of the protein molecule. We showed that along with the protein residues in the mutation sites, the residues in the cleavage sites are also involved in important interactions with the protein core. Moreover, the per residue importance profile shows the crucial role of the protein inner core in maintaining the stability of the protein. In addition to that, residues in the α1 strand are found to be important which can be experimentally validated for their influence on the protein stability and hence its pathogenesis. This information will help biomedical researchers working in emerging strategies towards TDP-43 disaggregation and develop prospective therapeutics in the future.

## 5. Declaration of competing Interests

The authors declare they have no conflict of interests.

## 6. Funding

This research did not receive any specific grant from funding agencies in the public, commercial, or not-for-profit sectors.

## Supplementary for

### The Random Forest (RF)

Let the feature dataset used for training is bootstrapped 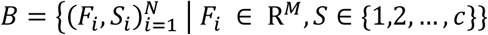, where *F_i_* are the feature set (variables) and *S* represents its respective label (the mutation state). Let *N* and *M* denotes the cardinality of samples used for training and the features respectively. For F be a given input instance and the prediction of the *K^th^* tree *T_k_* is represented as 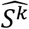. The prediction obtained by the RF as an ensemble of *K* is:

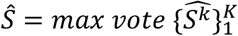

#### Pseudo-Code

The input be the bootstrapped training dataset *B*

|*mtry*| = *subspace size*,

|*F*| = *number of tress*

- For *k* → 1 *to K do*:

a. B_*k*_ samples are selected from the input to produce bootstrapped *B*
b. *mtry* features are chosen at random.
c. For *m* → 1 *to* ||mtry|| do
d. The amount of decrease in the node impurity (Gini Impurity) is calculate
e. Most contributing variable in the impurity decline is chosen and the node is then split into two daughter/child nodes
- The ensemble of the *K* trees produce a RF

While bootstrapping the feature dataset, due to the sampling with replacement not all the samples were used to prune the tree. These samples are called the *in-bag* samples. The left-out instances are coined as *out-of-bag* (OOB) sample. The OOB samples are exploited to calculate the error in prediction for each generated random forest also called as the OOB error rate.

The OOB value is given as: 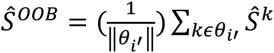, where 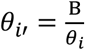, i’ and *i* denotes the out-of-bag and in-bag sampled instances, ||*θ_i_*|| is the number of OOB instances. The OOB prediction error is:

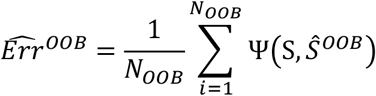

Here Ψ(·) is the error function and *N_OOB_* is OOB sample’s size.

### Gradient Boosting Algorithm

Boosting algorithms are gravitating a lot of attention in recent years. Of this, gradient boosting algorithm (GBM) are classification and regression models, that generates a prediction model through an ensemble of weak decision tress. Although a tree-based model, GBM outperforms the random forest models in terms of accuracy and speed [1,2,3]. The GBM model is built in a stage-wise fashion while introducing an arbitrary differentiable loss function [4,5]. Friedman introduced GBM models for regression and from there on the regularization and generalization of GBM emerged [6,7].

Here we state the pseudocode of a GBM machine. A preeminent view of GBM can be found in Friedman et al. [5].

Let the Data set be *X* as 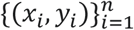. Let the differentiable loss function be defined as *L*(*y*, *F*(*x*)) and the number of iterations be *M*

The algorithm then follows:

- he model is initialized as constant value. The constant here is basically the mean target value.

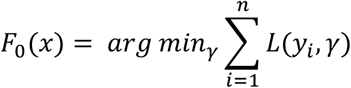
- For *m* = 1 *to M*:

- The pseudo-residuals are calculated as:

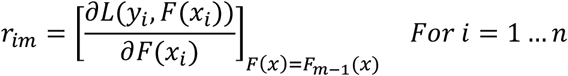
- The weak learner is fitted closed under the scaling of the calculated pseudo-residuals *h_m_*(*x*). This mean that the training is carried out over the new derived data set 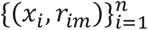.
- By solving the optimization problem, the *γ_m_* is calculates:

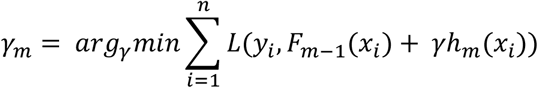
- The model is the updated

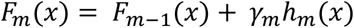
- Output *F_M_*(*x*)

### CV Table

**Table S1.** The classification performance metric of all the models for each type of mutation type. Accuracy = TP + TN/(TP+TN+FP+FN), Sensitivity =TP/(TP+FN), Specificity =TN/(TN+FP) and Precision = TP/(TP+FP) where TP, TN, FP and FN are True Positive, True Negative, False positive and False Negative classifications respectively.

| Model | Mutation Type | Accuracy | Sensitivity | Specificity | Precision |
| --- | --- | --- | --- | --- | --- |
| Random Forest | D168G | 0.999 | 0.998 | 1.000 | 1.000 |
|  | D168G_I169A | 1.000 | 1.000 | 1.000 | 1.000 |
|  | I169A | 0.996 | 0.992 | 1.000 | 1.000 |
| Elastic Net | D168G | 1.000 | 1.000 | 1.000 | 1.000 |
|  | D168G_I169A | 1.000 | 1.000 | 1.000 | 1.000 |
|  | I169A | 0.995 | 0.990 | 1.000 | 1.000 |
| LASSO | D168G | 1.000 | 1.000 | 1.000 | 1.000 |
|  | D168G_I169A | 1.000 | 1.000 | 1.000 | 1.000 |
|  | I169A | 0.999 | 0.998 | 1.000 | 1.000 |
| XGBoost | D168G | 1.000 | 1.000 | 1.000 | 1.000 |
|  | D168G_I169A | 1.000 | 1.000 | 1.000 | 1.000 |
|  | I169A | 0.997 | 0.995 | 1.000 | 1.000 |

### Hyperparameter Table

**Table S2.** The hyperparameters employed for the machine learning models.

| Model | Critical Parameters | Note |
| --- | --- | --- |
| LASSO | method = "glmnet",<br>lambda= seq(0.0001, 1, length = 5) | The alpha is not declared, setting 0 by default thus performing LASSO |
| RandomForest | ntree=300,<br>mtry= 8 | ntree employed here is the default value which was tested against 250, 300 and 400. |
| Elastic Net | method = "glmnet",<br>alpha= seq(0,1,length=10),<br>lambda= seq(0.0001, 1, length = 5) | Both alpha and beta were searched within the given possible set |
| XGBoost | Objective = Multi:softprob,<br>eval_metric = mlogloss,<br>num_class = 4,<br>nrounds = 1000,<br>eta = 0.01,<br>max.depth = 3,<br>gamma=0,<br>subsample=1 | The other parameters were kept in default |

**Fig. S1.**
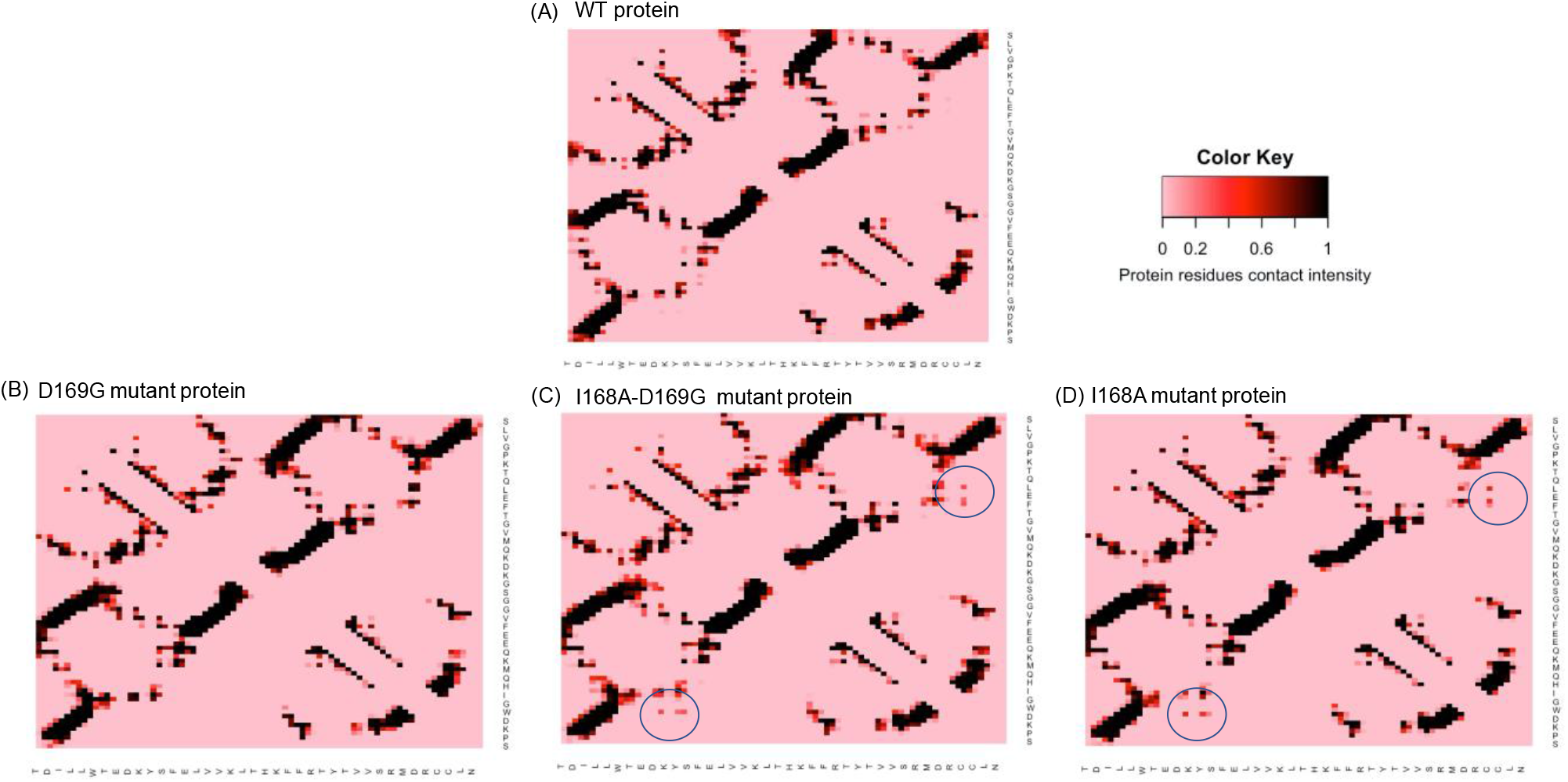
Contact map for (A) Wild-type (B) D169G (C) I168A-D169G (D) I168A proteins calculated throughout the simulation time. The darker the spot in the contact map, the longer the contact was formed between the pair of residues.

**Fig. S2.**
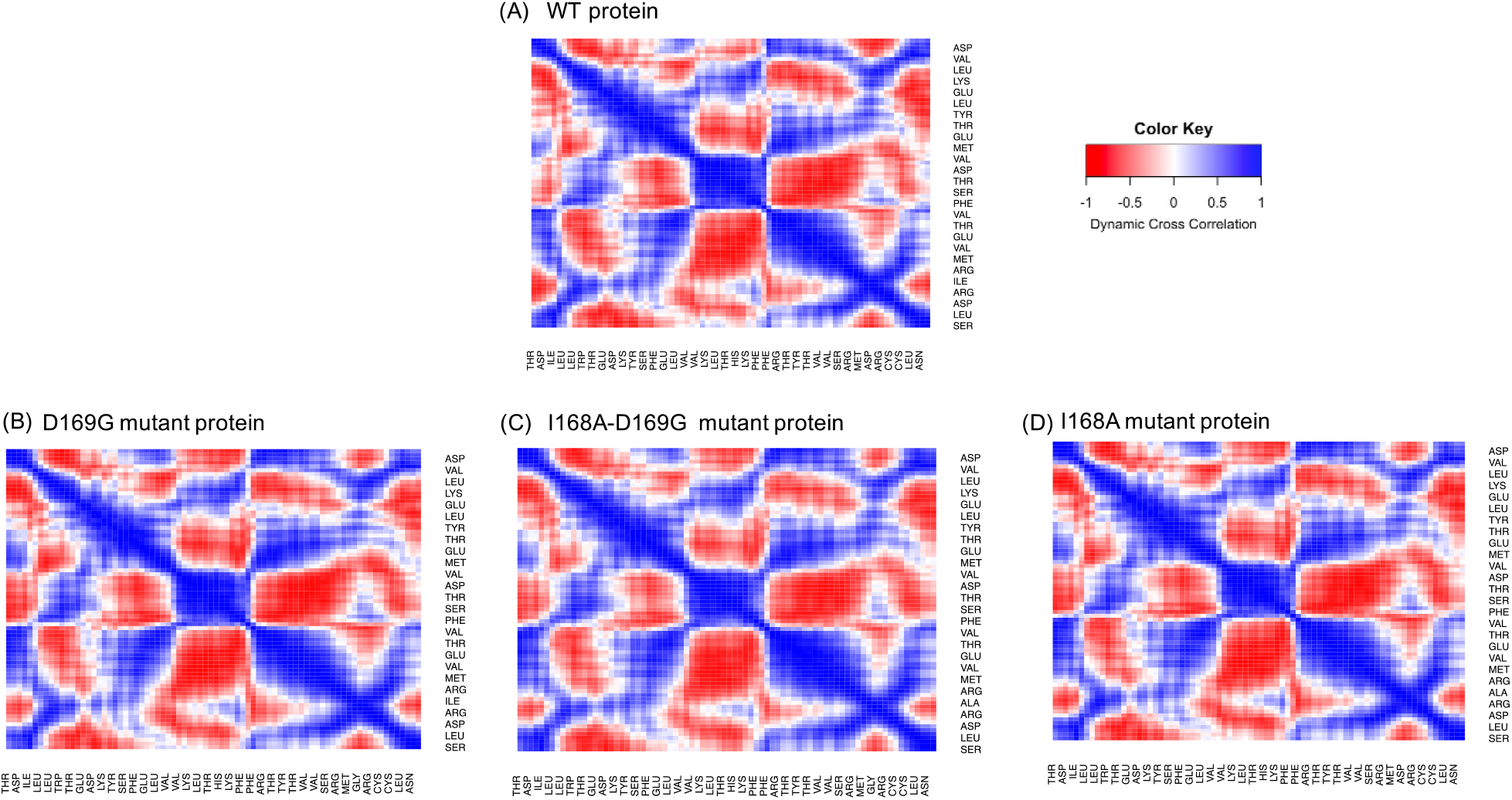
Dynamic cross-correlation map of (A) Wild-type (B) D169G (C) I168A-D169G (D) I168A proteins throughout the simulation time. The negative value represents anti-correlated motion whereas the positive value represents correlated motion.

**Fig. S3.**
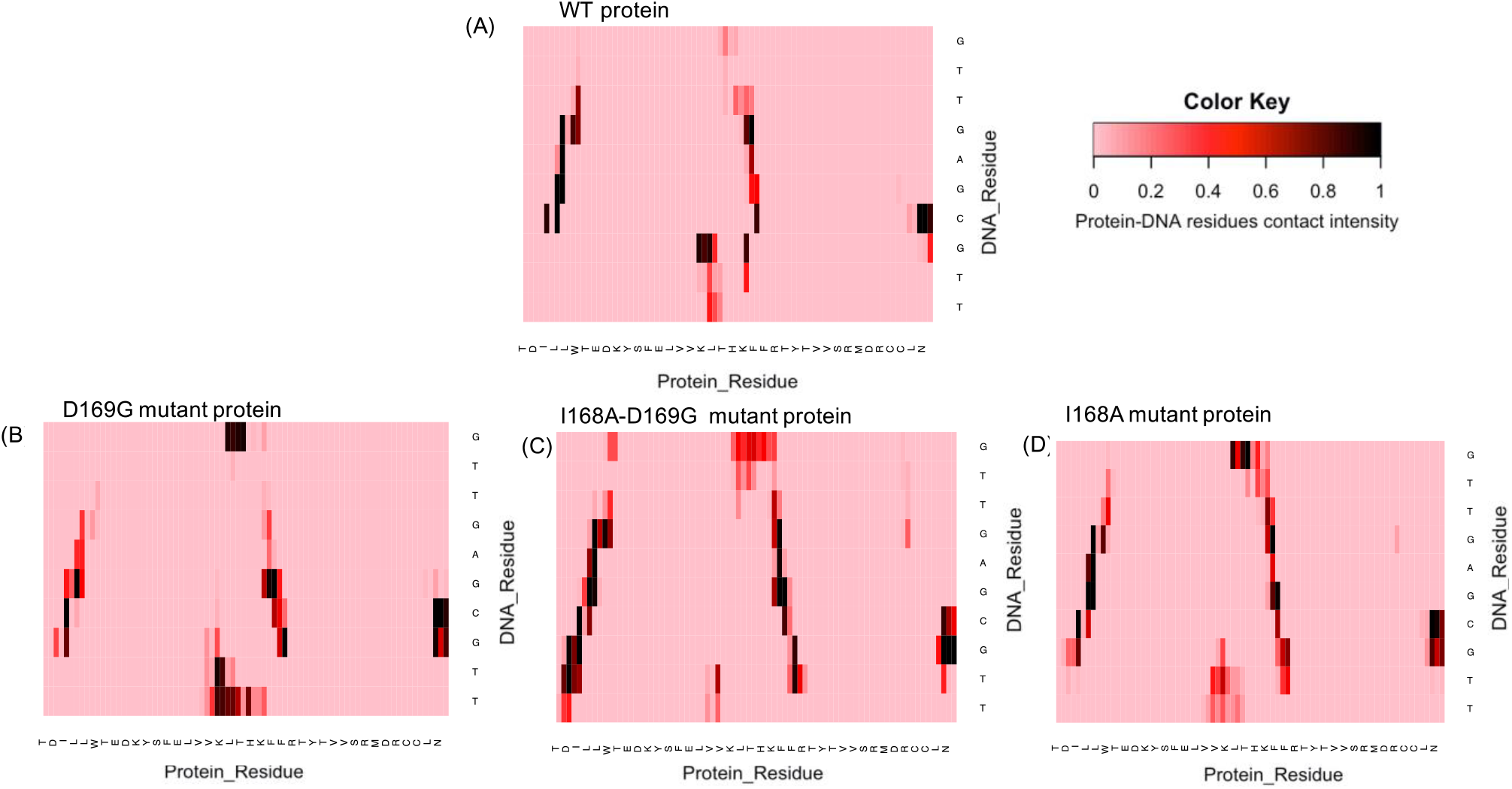
Protein-DNA contact map for (A) Wild-type (B) D169G (C) I168A-D169G (D) I168A proteins throughout the simulation time. The darker the spot in the contact map, the longer the contact was formed between the pair of residues.

### ROC AUC

**Fig. S4.**
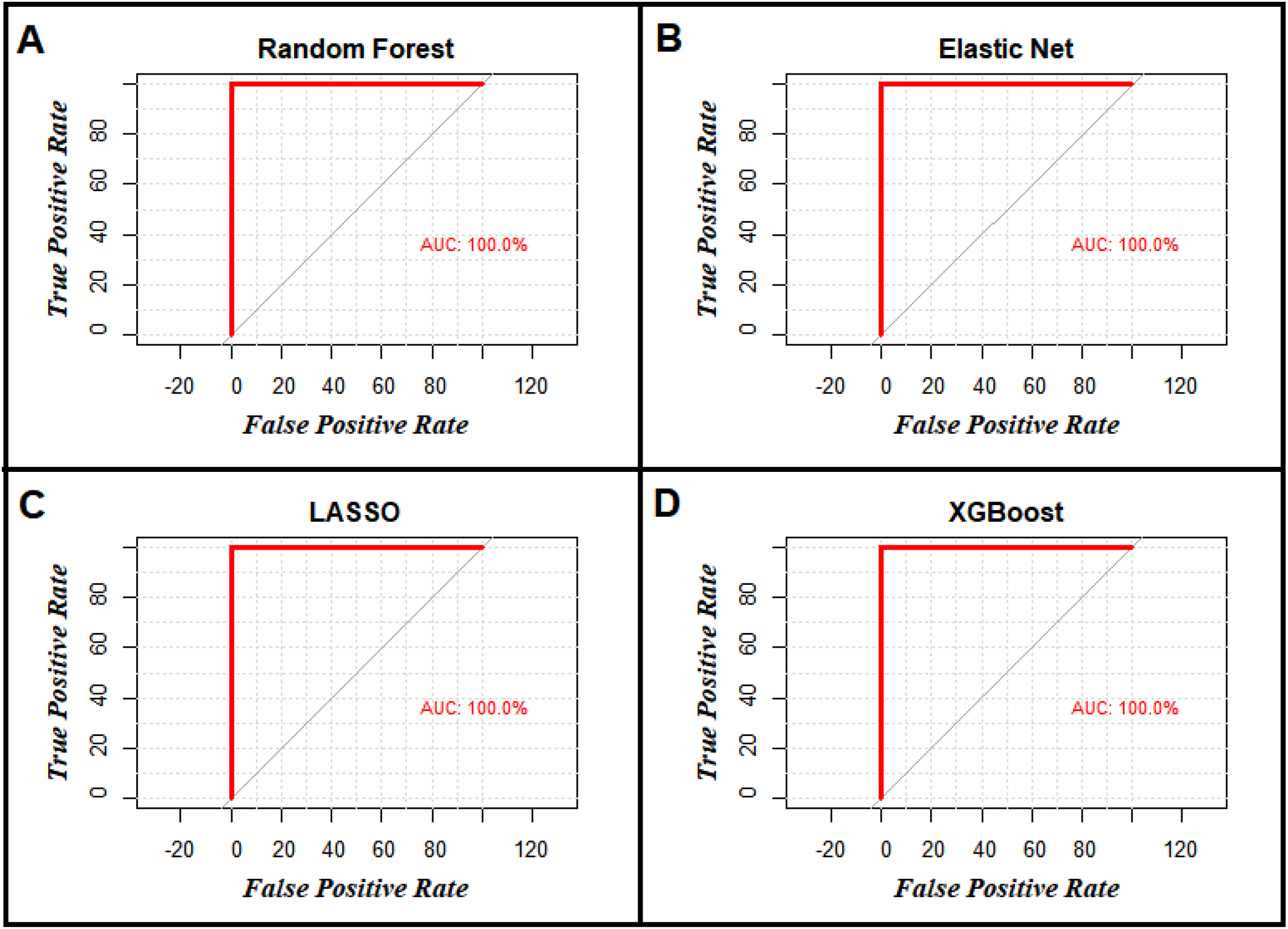
The ROC curve for each type of model (A) RandomForest, (B) Elastic Net, (C) LASSO, (D) XGBoost; plotted for the five-fold test dataset. The AUC (Area Under Curve) is observed to be 100% for all models when trained for all labelled mutated states.

**Fig. S5.**
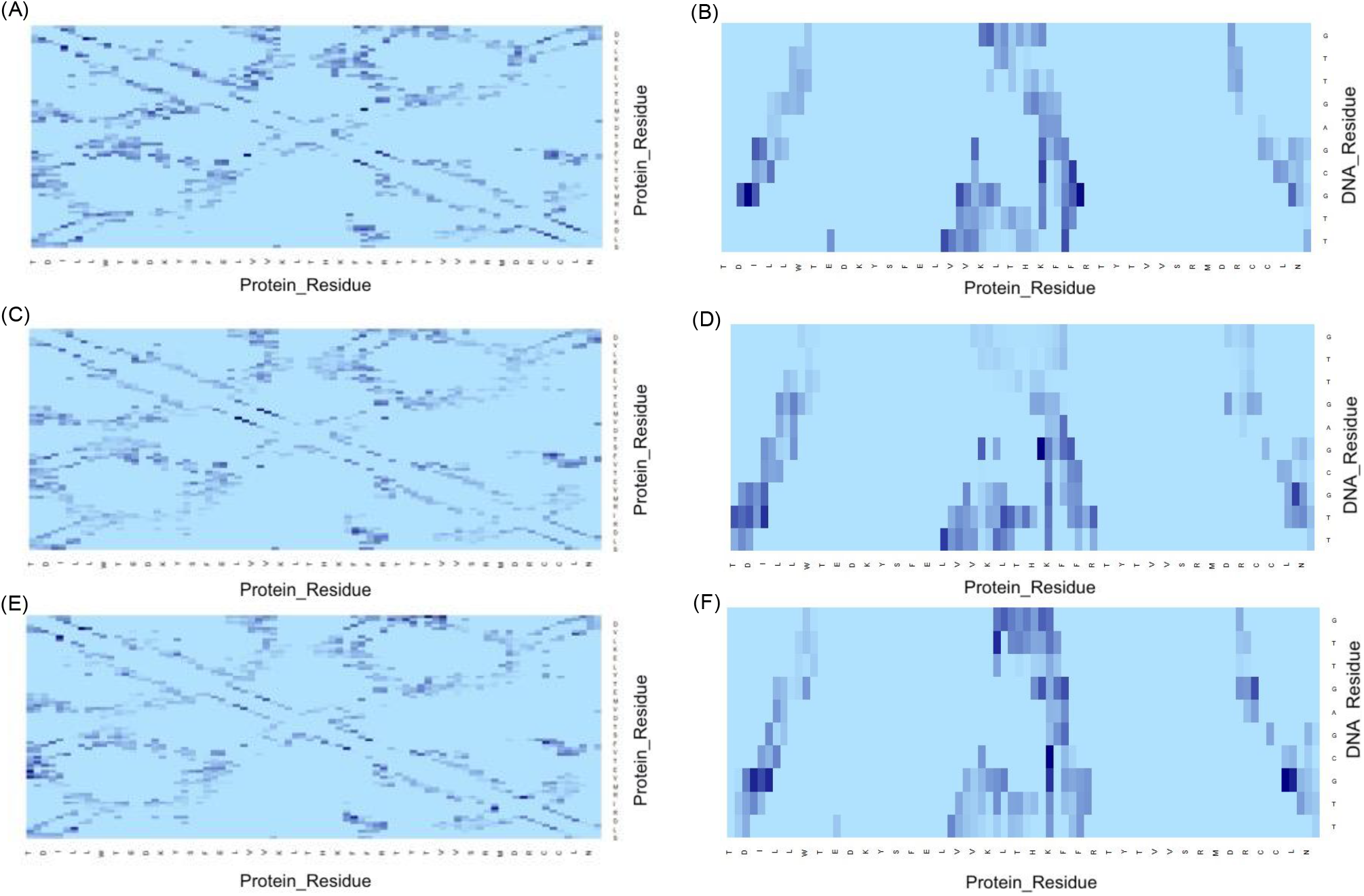
Importance Interaction map of Protein-protein interactions (A,C,E) and protein-DNA interactions(B, D, F) for (A,B) D169G (C,D) I168A-D169G (E,F) I168A mutant proteins. The colour intensity shows the importance of the interaction where the darker shade represents higher importance. For clear representation, ever third residue in the protein sequence are labelled in the plots.

**Fig. S6.**
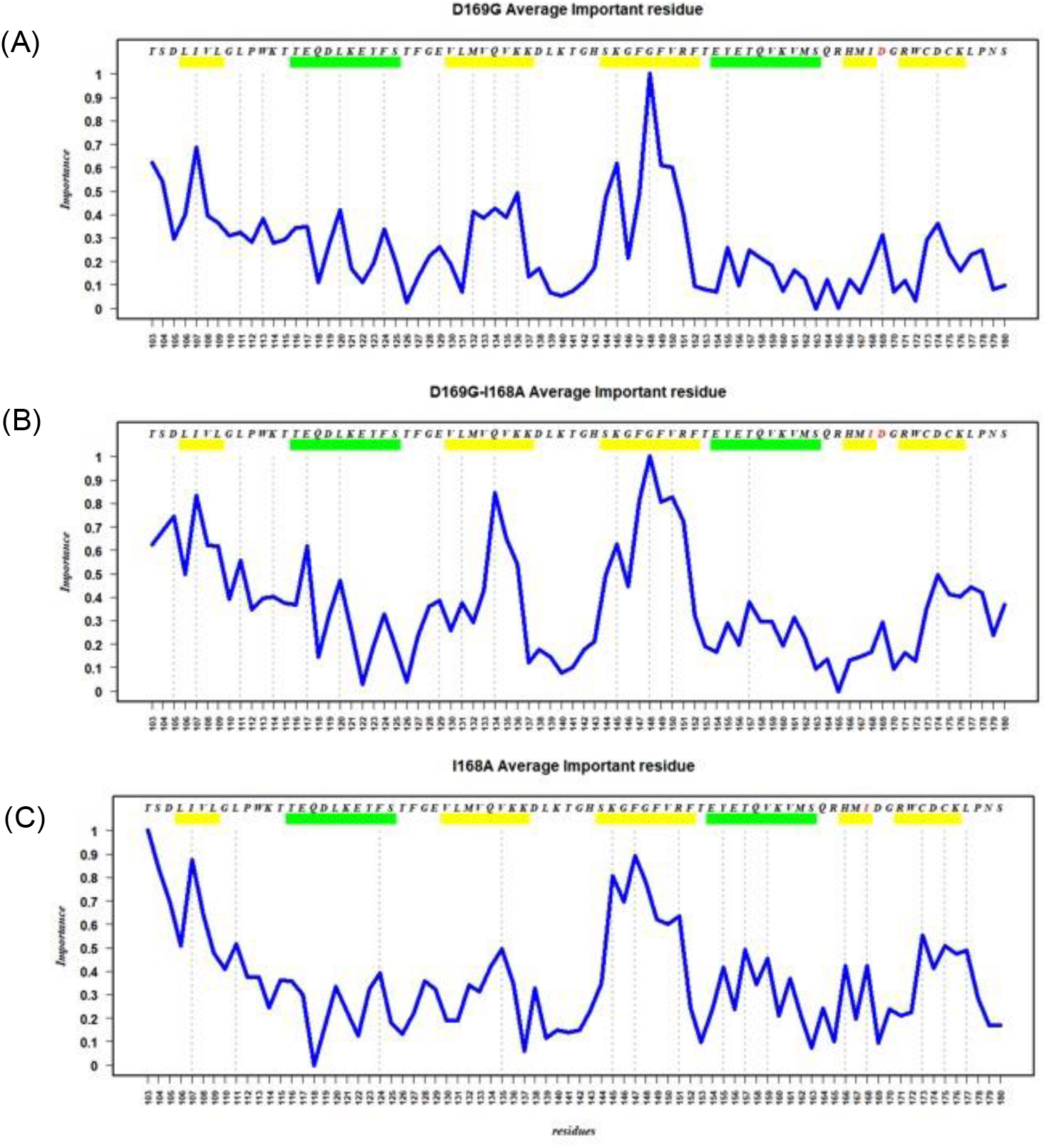
Important residue map of all the residues in the RRM1 motif for (A) D169G (B) I168A-D169G (C) I168A proteins. The green and yellow bars represent the alpha helix and beta strand respectively.

## Notes

### Competing Interest Statement

The authors have declared no competing interest.

